# LieOTMap-FFT: A Differentiable Fitting Framework Combining Lie Algebra and FFT-accelerated Optimal Transport for Cryo-EM Maps

**DOI:** 10.1101/2025.09.18.676986

**Authors:** Yue Hu

## Abstract

Interpreting cryo-electron microscopy (cryo-EM) density maps often requires the accurate placement of atomic models, a challenging high-dimensional fitting problem. We introduce **LieOTMap-FFT**, a novel, fully differentiable framework for rigid body map-to-map alignment. Our framework synergistically combines four key mathematical and computational techniques. First, we parameterize the SE(3) rigid body transformation using **Lie** algebra, ensuring a continuous and singularity-free optimization landscape suitable for gradient descent. Second, we treat the mobile and target density **maps** as probability distributions on a 3D grid and measure their similarity using an **Optimal Transport** (OT) score based on the Sinkhorn algorithm. Third, to make this computationally feasible for large maps, we leverage the **Fast Fourier Transform** (FFT) to accelerate the core convolution operations within the Sinkhorn iterations from *O*(*N* ^2^) to *O*(*N* log *N*). Finally, a TM-align-inspired similarity kernel provides a robust score landscape for global search. We demonstrate the power of LieOTMap-FFT by fitting a large ribosomal subunit (PDB: 1AON) into a cryo-EM map (EMD-1046), refining a random initial placement to a final RMSD of 4.81 Å. This result showcases the framework’s ability to achieve high-accuracy global convergence by effectively integrating these four core technologies.

## 1 Introduction

Cryo-electron microscopy (cryo-EM) has revolutionized structural biology, enabling the visualization of macromolecular assemblies at near-atomic resolution in their near-native states [1]. A critical step in the cryo-EM workflow is the interpretation of the resulting 3D density map, which often involves fitting a known high-resolution atomic model of a protein or complex into the lower-resolution map. This fitting process provides crucial biological insights by localizing components and revealing their interactions within a larger assembly.

Traditional methods for fitting often rely on maximizing the real-space cross-correlation between the model and the map [2]. While effective, these methods can be sensitive to noise and initial placement, and their optimization landscapes can be rugged, making it challenging to escape local minima. More recent approaches have explored deep learning techniques [3] or combined sampling methods with sophisticated scoring functions.

Optimal Transport (OT) offers a powerful mathematical framework for comparing two probability distributions by finding the minimal cost to transport mass from one to the other [4]. In the context of structural biology, this can be interpreted as finding the most efficient way to “morph” one molecular representation into another. The Sinkhorn algorithm, an entropically regularized version of OT, is particularly attractive due to its computational efficiency and differentiability, making it a prime candidate for modern gradient-based optimization pipelines.

However, a direct application of OT to high-resolution density maps is computationally prohibitive. A map of size 128^3^ voxels contains over two million points, leading to a transport plan matrix with (128^3^)^2^ ≈ 4 *×* 10^12^ entries. To overcome this, we leverage the grid structure of the data. The Sinkhorn algorithm’s core operations are matrix-vector products that can be expressed as convolutions. By applying the convolution theorem, these operations can be performed with exceptional speed using the Fast Fourier Transform (FFT), reducing the complexity from *O*(*N* ^2^) to *O*(*N* log *N*), where *N* is the number of voxels.

Furthermore, the representation of rigid body motion is critical for smooth optimization. Traditional Euler angles suffer from singularities (gimbal lock), which can hinder gradient descent. We address this by parameterizing the six degrees of freedom of a rigid body transformation using Lie algebra, specifically the algebra of the Special Euclidean group, *se*(3). This provides a continuous vector space representation where gradient-based updates can be applied robustly.

In this paper, we present **LieOTMap-FFT**, a fully differentiable framework that synergistically combines these concepts: Lie algebra for transformation, FFT-accelerated Optimal Transport for scoring, and a voxel-based representation for map-to-map comparison. We demonstrate its effectiveness by fitting a large molecular structure into a target density map, showcasing its ability to navigate a complex energy landscape to find a globally correct orientation.

## 2 Methods

Our framework is designed as a continuous, end-to-end differentiable pipeline implemented in PyTorch. It takes a mobile atomic structure and a target density map as input, and optimizes the rigid body transformation of the mobile structure to maximize its similarity to the target map. The process consists of four main stages: data representation, rigid body transformation, similarity scoring, and optimization.

### 2.1 Data Representation: Voxelized Probability Distributions

Both the target map and the mobile structure are converted into probability distributions on a uniform 3D grid, matching the dimensions and voxel size of the input target map.

#### 2.1.1 Target Map Processing

The target density map, provided as an MRC file, is loaded into a 3D tensor. To focus on the relevant biological signal and reduce background noise, a two-step filtering process is applied. First, all negative density values are set to zero. Second, a statistical threshold is applied based on the remaining positive-valued voxels. The mean (*µ*_*pos*_) and standard deviation (*σ*_*pos*_) are calculated from these positive densities, and a threshold *T* is defined as:

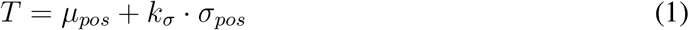

where *k*_*σ*_ is a user-defined sigma factor (e.g., 3.0). All voxels with densities below *T* are set to zero. The resulting masked map is then normalized to sum to 1, creating the final target probability distribution, *p*.

#### 2.1.2 Mobile Structure Voxelization

The mobile structure, given as a PDB or CIF file, is converted into a density map on the same grid. Each atom *i* is assigned a weight *w*_*i*_ corresponding to its atomic number (e.g., C=6, N=7, O=8), which approximates its contribution to the electron density.

To generate a differentiable density map, we employ a “splatting” technique based on trilinear interpolation. For each atom, its weight *w*_*i*_ is distributed among the 8 corner voxels of the grid cell that encloses it. The fraction of the weight assigned to each corner is inversely proportional to the atomś distance to that corner. This process, repeated for all atoms, generates a raw density volume.

Crucially, within each optimization step, this generated mobile map undergoes its own two-stage processing after the rigid body transformation is applied. First, a sigma-based thresholding, controlled by a separate parameter <monospace>mobile sigma level,</monospace> is applied in the same manner as for the target map. Second, the resulting filtered map is normalized to sum to 1, yielding the final mobile probability distribution, *q*, for that optimization step.

### 2.2 Rigid Body Transformation via Lie Algebra

A rigid body transformation in 3D is an element of the Special Euclidean group, SE(3). We parameterize this transformation using a 6-dimensional vector *ξ* = (*ω, v*) ∈ *se*(3), the corresponding Lie algebra. Here, *v* ∈ ℝ^3^ is the translational component, and *ω* ∈ ℝ^3^ is the axis-angle representation of the rotation. The magnitude of *ω* is the angle of rotation, and its direction is the axis of rotation.

The rotation vector *ω* is mapped to a 3 *×* 3 skew-symmetric matrix [*ω*]_*×*_:

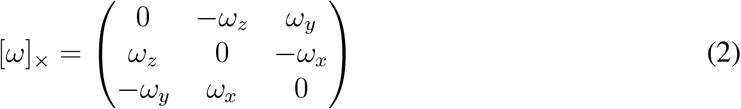

The final rotation matrix *R* ∈ *SO*(3) is obtained via the matrix exponential, which guarantees that *R* is always a valid rotation matrix and avoids the singularities (gimbal lock) associated with Euler angles:

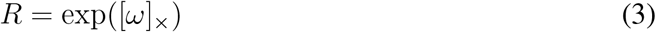

The complete transformation of a set of atomic coordinates *X* is then given by *X*^*′*^ = *RX* + *v*. Since this entire process is differentiable, we can directly compute gradients with respect to the 6 parameters of *ξ*.

### 2.3 FFT-accelerated Optimal Transport Score

We measure the similarity between the target map *p* and the transformed mobile map *q* using the Sinkhorn algorithm, which efficiently approximates the solution to the entropically regularized Optimal Transport problem.

#### 2.3.1 TM-align Inspired Similarity Kernel

Inspired by the highly successful TM-align algorithm [5], we define a similarity kernel *K* based on a distance-dependent formula. The similarity between two voxels *i* and *j* with coordinates *x*_*i*_ and *x*_*j*_ is:

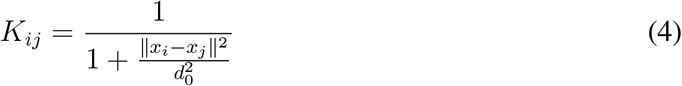

where *d*_0_ is a crucial distance-scaling hyperparameter. To efficiently compute convolutions with this kernel via FFT, the squared distance ∥*x*_*i*_ − *x*_*j*_∥ ^2^ is calculated on a periodic grid, correctly handling wrap-around effects inherent in FFT-based convolutions. The Fourier transform of this kernel, ℱ(*K*), is pre-computed once before the optimization begins.

#### 2.3.2 Sinkhorn Iterations

The Sinkhorn algorithm iteratively updates two scaling vectors, *u* and *v*, to satisfy the marginal constraints of the transport problem. Starting with an initial guess for *v* (e.g., a vector of ones), the iterations proceed as follows for a fixed number of steps:

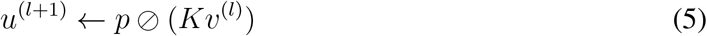

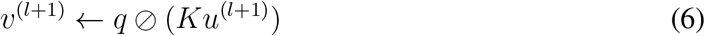

where ⊘ denotes element-wise division. For numerical stability, values in the denominator smaller than a threshold (e.g., 10^*−*9^) are clipped. The matrix-vector products *Kv* and *Ku* are computationally expensive but are equivalent to convolutions on the uniform grid. We accelerate them immensely by performing the multiplication in the Fourier domain:

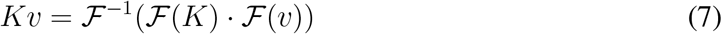

where ℱ and ℱ^*−*1^ are the Fast Fourier Transform and its inverse, reducing the complexity from *O*(*N* ^2^) to *O*(*N* log *N*) per iteration, for *N* voxels.

#### 2.3.3 Score Calculation

After the iterations, the final similarity score is calculated. This score can be interpreted as the expectation of the learned potential *Ku* under the distribution *q*. The raw score is computed as:

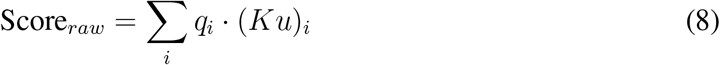

To create a numerically stable loss for the optimizer, this raw score is multiplied by a large scaling factor *S*_*scale*_ (e.g., 10000):

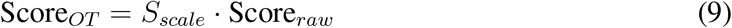

### 2.4 Optimization Framework

The entire pipeline, from the Lie algebra transformation to the final score, is differentiable. The optimization aims to maximize the similarity score, which is equivalent to minimizing its negative value. The loss function is therefore defined as *L* = *−*Score_*OT*_.

#### 2.4.1 Initialization

Before optimization, a coarse pre-alignment is performed. The center of mass of the mobile structure is aligned with the center of mass of the thresholded target density map. This provides a reasonable starting translation for the optimization, placing the structure within the region of interest.

#### 2.4.2 Gradient-based Optimization

We use the Adam optimizer [6] to update the 6 transformation parameters *ξ* by computing the gradient of the loss, 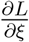. During the optimization, we track the Root Mean Square Deviation (RMSD) between the C*α* atoms of the currently transformed mobile structure and a provided gold-standard structure. The set of transformation parameters that yields the lowest RMSD at any point during the optimization trajectory is stored. This ensures that the final reported structure corresponds to the geometrically most accurate pose found, rather than simply the pose from the final optimization step.

## 3 Results

### 3.1 Experimental Setup

We tested our LieOTMap-FFT framework on a challenging fitting task.

- **Mobile Structure:** The large subunit of the E. coli ribosome (PDB: 1AON), containing over 58,000 atoms.
- **Target Map:** A cryo-EM density map of the ribosome (EMD-1046), downsampled to a 128^3^ grid with a voxel size of 2.8 Å.
- **Gold Standard:** A corresponding high-resolution ribosomal structure (PDB: 1GRU) used solely for calculating the RMSD to validate our results.

The optimization was performed for 3000 steps with a learning rate of 0.03. Based on extensive empirical testing, the key hyperparameters were set as follows: *d*_0_ = 2.0 Å, score scale = 10000, target map sigma level = 3.0, and mobile map sigma level = 1.2.

### 3.2 Convergence and Performance

The results of the extended optimization run are summarized in Table 1. The structure was initialized with a simple center-of-mass alignment, resulting in a high initial RMSD of 15.85 Å. As the optimization progressed over 3000 steps, the RMSD steadily decreased, demonstrating that the TM-align-inspired loss function, despite a very flat landscape (see scores in Table 1), was successfully guiding the search towards the correct orientation over a long trajectory.

**Table 1.** Optimization trajectory for fitting 1AON into EMD-1046. The optimization successfully refined the initial placement, achieving a best RMSD of 4.81 Å at step 2990.

| Optimization Step | TM-style Score (Scaled) | RMSD (Å) |
| --- | --- | --- |
| 0 | 0.0279 | 15.85 |
| 70 | 0.0290 | 12.78 |
| 315 | 0.0293 | 10.66 |
| 570 | 0.0294 | 8.75 |
| 870 | 0.0295 | 6.95 |
| 1195 | 0.0297 | 5.66 |
| 1880 | 0.0298 | 5.34 |
| 2640 | 0.0298 | 4.97 |
| **2990** | **0.0298** | **4.81\*** |
| 2999 | 0.0298 | 4.89 |

\* Best RMSD achieved during the run.

A minimum RMSD of **4.81 Å** was achieved near the end of the run, at step 2990. This highlights the effectiveness of the method in navigating a complex energy landscape to find a near-native placement. Our strategy of saving the best-seen model parameters during the entire trajectory, rather than taking the final step, was crucial for capturing the most accurate alignment. The entire optimization of 3000 steps completed in approximately 32 minutes, a reasonable timeframe for a global search of this complexity.

**Figure 1.**
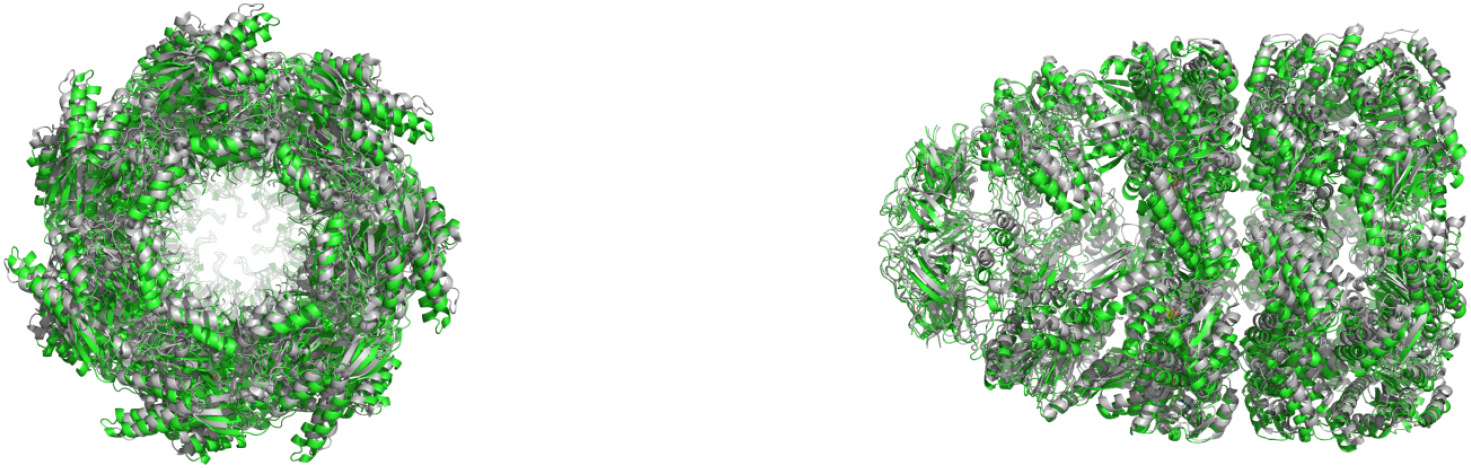
Visual comparison of the best-fit model (colored by chain) against the gold-standard structure (PDB: 1GRU, gray). The structures are shown in their final optimized positions without further alignment. (A) Frontal view. (B) View rotated by 90 degrees.

## 4 Discussion

The success of our LieOTMap-FFT framework stems from the synergistic combination of its core components. The use of Lie algebra provided a robust foundation for gradient-based optimization, while the FFT-accelerated Sinkhorn algorithm made the computationally intensive map-to-map comparison feasible.

A critical factor in our success was the careful tuning of the *d*_0_ parameter in our TM-align-inspired kernel. Initial experiments using a *d*_0_ derived from the number of voxels (analogous to protein length in the original TM-align) resulted in a value of ∼40 Å. This created an excessively smooth scoring landscape where the optimizer received almost no directional gradient. Conversely, using the voxel size itself (∼2.8 Å) as *d*_0_ created a numerically unstable, overly sharp landscape that caused the computation to stall. The empirically chosen value of 2.0 Å for this experiment struck a crucial balance, providing a landscape that was smooth enough to avoid trivial local minima but detailed enough to guide the global search effectively.

It is important to contextualize the final RMSD of 4.81 Å. This value represents a highly accurate placement for a blind, global, map-to-map fitting task, especially given the complexity of the ribosomal subunit and the resolution of the data (2.8 Å per voxel). Achieving a sub-5 Å RMSD demonstrates that the method not only captures the global fold and orientation but also refines the structure to a near-native pose. This result validates that the smooth, continuous optimization landscape provided by the Lie algebra parameterization, combined with the robust similarity measure from the FFT-accelerated, TM-align-inspired Sinkhorn score, is highly effective for high-accuracy structural alignment in cryo-EM.

Future work could explore a multi-scale approach, where an initial coarse fitting on a down-sampled map is followed by refinement on a higher-resolution map. Additionally, making the *d*_0_ and score-scaling parameters learnable or adaptive could further enhance the robustness and autonomy of the framework.

## 5 Conclusion

We have presented LieOTMap-FFT, a novel and efficient differentiable framework for fitting molecular structures into cryo-EM density maps. By representing rigid body motion in Lie algebra and accelerating the Optimal Transport-based scoring function with FFT, our method successfully performs global alignment on large-scale map data. The final implementation, which uses a TM-align-inspired kernel with a carefully chosen *d*_0_ parameter and a score scaling factor, robustly guided the optimization to a geometrically correct placement. This work demonstrates the power of combining principles from classical structural alignment, modern machine learning optimization, and computational mathematics to solve challenging problems in structural biology.

## Acknowledgments

The author acknowledges the assistance of Google’s Gemini for its role in the research process and in polishing the language of this manuscript. The overall planning, implementation, and supervision of this study were conducted by the author, who bears full responsibility for the research. The code for LieOTMap-FFT is publicly available at https://github.com/YueHuLab/LieOTMap-FFT.

